# Genome-scale metabolic modelling enables deciphering ethanol metabolism via the acrylate pathway in the propionate-producer Anaerotignum neopropionicum

**DOI:** 10.1101/2022.04.11.487872

**Authors:** Sara Benito-Vaquerizo, Ivette Parera Olm, Thijs de Vroet, Peter J. Schaap, Diana Z Sousa, Vitor A.P. Martins dos Santos, Maria Suarez-Diez

## Abstract

**Background:** Microbial production of propionate from diluted streams of ethanol (e.g., deriving from syngas fermentation) is a sustainable alternative to the petrochemical production route. Yet, few ethanol-fermenting propionigenic bacteria are known, and understanding of their metabolism is limited. *Anaerotignum neopropionicum* is a propionate-producing bacterium that uses the acrylate pathway to ferment ethanol and CO_2_ to propionate and acetate. In this work, we used computational and experimental methods to study the metabolism of *A. neopropionicum* and, in particular, the pathway for conversion of ethanol into propionate.

**Results:** Our work describes iANEO_SB607, the first genome-scale metabolic model (GEM) of *A. neopropionicum*. The model was built combining the use of automatic tools with an extensive manual curation process, and it was validated with experimental data from this and published studies. The model predicted growth of *A. neopropionicum* on ethanol, lactate, sugars and amino acids, matching observed phenotypes. In addition, the model was used to implement a dynamic flux balance analysis (dFBA) approach that accurately predicted the fermentation profile of *A. neopropionicum* during batch growth on ethanol. A systematic analysis of the metabolism of *A. neopropionicum* combined with model simulations shed light into the mechanism of ethanol fermentation via the acrylate pathway, and revealed the presence of the electron-transferring complexes NADH-dependent reduced ferredoxin:NADP^+^ oxidoreductase (Nfn) and acryloyl-CoA reductase-EtfAB, identified for the first time in this bacterium.

**Conclusions:** The realisation of the GEM iANEO_SB607 is a stepping stone towards the understanding of the metabolism of the propionate-producer *A. neopropionicum*. With it, we have gained insight into the functioning of the acrylate pathway and energetic aspects of the cell, with focus on the fermentation of ethanol. Overall, this study provides a basis to further exploit the potential of propionigenic bacteria as microbial cell factories.

## Background

Propionic acid is a naturally-occurring carboxylic acid produced by propionigenic bacteria as end-product of their anaerobic metabolism. It is an important intermediate in anaerobic fermentative processes such as those occurring in the human gut, anaerobic digesters and cheese production. It is also an essential platform chemical in the manufacture of cellulose-derived plastics, cosmetics and pharmaceuticals and, due to its antimicrobial properties, it can be used as food preservative [1, 2]. At present, industrial production of propionic acid is based on petrochemical processes, but efforts are being made to develop sustainable production platforms based on the use of propionigenic bacteria as biocatalysts [1, 2]. Microbial production of propionic acid has been researched for over 150 years, however industrial implementation is still limited mainly due to low productivities, which render such processes economically noncompetitive [1, 2, 3]. So far, most approaches have considered strains of the genus *Propionibacterium* - well-studied due to their involvement in cheese production [2] -, and have focused on the use of sugars as feedstock. However, the chemical industry is increasingly required to rely on the use of non-conventional, inexpensive raw materials to minimize its carbon footprint [4]. Ethanol, a low-priced common end-product of many fermentations, is regarded as one of such feedstocks [4, 5]. Moreover, ethanol can be synthesised from CO, CO_2_ and H_2_ (syngas) by acetogenic bacteria. Syngas-to-ethanol fermentation technology has been deployed at large scale, and recent advances are expected to accelerate its development in the years to come [6, 7, 8].

*Anaerotignum neopropionicum*, formerly *Clostridium neopropionicum* [9], was the first representative of the ethanol-fermenting, propionate-producing bacteria. It was isolated in 1982 from an anaerobic digester treating wastewater from vegetable cannery [10]. The ability of converting ethanol to propionate is shared with only three other microbial species: the closest relative *Anaerotignum propionicum* [11] (formerly, *Clostridium propionicum* [9]), the sulphate-reducing bacterium *Desulfobulbus propionicus* [12, 13], and *Pelobacter propionicus* [14]. In these four microorganisms, ethanol oxidation to propionate occurs in the presence of CO_2_ with concomitant production of acetate, according to the theoretical Eq. (1). This ability of propionigenic bacteria could be exploited to upgrade dilute ethanol streams from beer production or syngas fermentation, among others. For example, Moreira et al. showed that co-cultures of acetogens and ethanol-consuming propionigenic bacteria can convert syngas into propionate [15]. In their study, the acetogen *Acetobacterium wieringae* was co-cultivated with *A. neopropionicum*; *A. wieringae* converted CO to ethanol, which was used by *A. neopropionicum* to produce propionate.

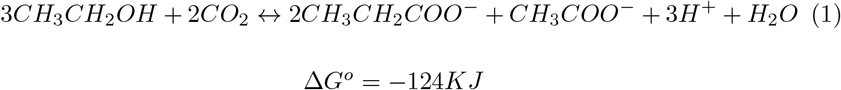

Two main pathways have been described for the fermentative production of propionic acid in bacteria: the methylmalonyl-CoA (also termed succinate pathway or Wood-Werkman cycle) and the acrylate pathway [1, 16]. Most of the known propionigenic bacteria, including strains of the genera *Propionibacterium* and *Cutibacterium*, use the methylmalonyl-CoA pathway for growth. The acrylate pathway is mostly found within members of the phylum Firmicutes [16]. Sugars and lactate are common substrates for these pathways. Ethanol fermenters *D. propionicus* and *P. propionicus* use the methylmalonyl-CoA pathway [13, 14], whereas *A. neopropionicum* and *A. propionicum* use the acrylate pathway [17].

To fully exploit the potential of microorganisms for biotechnological applications, it is fundamental to understand their metabolism and cellular processes. Genome-scale metabolic models (GEMs) and their analysis with COnstraint-Based Reconstruction and Analysis (COBRA) methods [18] have become indispensable tools in this regard [19, 20]. Flux balance analysis (FBA) is often used as the mathematical approach to explore the intracellular fluxes of GEMs under steady-state conditions (e.g., in chemostat cultivations) [21]. FBA can be extended to dynamic FBA (dFBA), which simulates the time-step evolution of individual steady-states that take place in time-varying processes, such as batch and fed-batch cultures [22]. A wide range of GEMs have been successfully implemented to unravel novel metabolic features of microorganisms, guide experimental design or improve bioprocess operation in mono- and co-cultivation. For instance, the reconstruction of the first GEM of *Clostridium ljungdahlii* (iHN637) demonstrated the essential role of flavin-based electron bifurcation in energy conservation during autotrophic growth [23]. FBA enabled the estimation of intracellular metabolic fluxes in the GEM of the acetogen *Clostridium autoethanogenum* (iCLAU786), helping to understand the effects of CO supplementation on CO_2_/H_2_-growing cultures [24]. A multi-species GEM was recently developed that described a syngas-fermenting co-culture composed of *C. autoethanogenum* and *Clostridium kluyveri*; the model provided valuable insight into the microbial interactions between the two microorganisms and predicted strategies for enhanced production of the end products butyrate and hexanoate [25].

Many propionigenic bacteria have been sequenced to date [26, 27, 28, 29, 30, 31], including the ethanol fermenters *D. propionicus* [32], *P. propionicus* [33], *A. propionicum* [29] and *A. neopropionicum* [31]. This has enabled the reconstruction of GEMs of some of these species. All GEMs of propionigenic bacteria published to date concern strains that harbour the methylmalonyl-CoA pathway. One of these works described the reconstruction of five *Propionibacterium freudenreichii* species using pan-genome guided metabolic analysis [34]. Navone et. al used the *Propionibacterium* subsp. *shermanii* and the pan-*Propionibacterium* GEMs to guide genetic engineering strategies for increased propionic acid production [35]. Sun et. al developed a constrained-based GEM of *P. propionicus* and validated fermentative growth of this strain on ethanol [36].

Here we describe iANEO_SB607, the first GEM of *A. neopropionicum* and the first to model the acrylate pathway in a propionigenic microorganism. The model was reconstructed using automatic tools followed by an extensive manual curation, which led us to the identification of electron-transferring enzymes involved in the acrylate pathway, cofactor regeneration and energy conservation. In addition, a physiological characterisation of *A. neopropionicum* in batch cultures was performed to validate and complement the reconstruction of the model. FBA was used to assess growth phenotypes on several carbon sources, and dFBA was applied to simulate batch growth of *A. neopropionicum* on ethanol, and ethanol plus acetate. The combination of in-depth modelling and experimentation has enabled us to examine in detail the metabolism of ethanol fermentation in this bacterium and to address pre-existing ambiguities.

## Materials and methods

### Reconstruction of the GEM iANEO_SB607

The genome-scale metabolic network of *A. neopropionicum* was reconstructed in four main steps. First, the genome sequence of *A. neopropionicum* DSM 3847^T^ (GCA_001571775.1) [31] was retrieved from the European Nucleotide Archive in FASTA format and was annotated using RAST [37]. An additional re-annotation was carried out using eggNOG-mapper [38]. The annotation file can be found in the public Gitlab repository. The second step was the generation of the draft model using ModelSEED [39]. For this, the RAST annotation file was imported into ModelSEED and a Gram-positive template was chosen to reproduce growth on rich medium. The draft model was downloaded in table format and SBML format. The third step consisted on the manual curation and refinement of the draft model. Every reaction entry was analysed individually and modifications were made on the table format file. Specifically, (i) unbalanced reactions were corrected based on charged formulas with the corresponding addition/deletion of H^+^ or H_2_O molecules; (ii) reaction direction was adjusted using eQuilibrator [40]. Reactions were considered reversible if the change in Gibbs free energy was between -30 and 30 KJ mol^-1^ at standard conditions for reactants/products, pH 7.3 and ionic strength 0.1 M. In cases where eQuilibrator did not retrieve information for a specific reaction, reaction direction was adjusted based on information from MetaCyc [41] and BIGG [42] databases. (iii) EC numbers were corrected or inserted for every reaction based on information from KEGG [43] and MetaCyc [41]. (iv) The original genes in Patric format [44] were replaced by the locus tag format (’CLNEO_XXXXX’) found in Uniprot [45] and BRENDA [46] databases. The re-annotation file was used to identify potential gene(s) associated to reactions that lacked a gene in the original RAST annotation. (v) The final step consisted of gap-filling, where reactions were added or removed to reproduce known or observed phenotypes. Gap-filling was done combining a computational and a manual approach: an automatic gap-filling process was run using the KBase pipeline[47], while the manual curation was based on experimental data obtained in this study and published. The final model, iA-NEO_SB607, can be found in the git repository in Table format, json and SBML L3V1 [48] standardization. Furthermore, the different versions were combined in an OMEX archive file [49] deposited in BioModels [50] and assigned the identifier MODEL2201310001.

#### Generation of the biomass synthesis reaction and sensitivity analysis

The biomass reaction of *A. neopropionicum* was adapted from the biomass reactions of *Clostridium beijerinckii* (GEM iCM925 [51]) and *Clostridium autoethanogenum* (GEM iCLAU786 [52]). The composition of the main building blocks was maintained but, based on the protocol of Thiele and Palsson [53], protons were stoichiometrically added to the hydrolysis part of the biomass synthesis reaction. Protons were also added to the reactions of DNA, RNA, proteins, teichoic acids and peptidoglycans synthesis in line with the ATP associated to polimerization. The DNA composition was determined based on the GC content of the genome of *A. neopropionicum* and it was adjusted in the reaction associated to the biosynthesis of DNA. The fatty acids composition was adjusted based on reported experimental data for *A. neopropionicum* [9].

A sensitivity analysis was performed by modifying the content of proteins, phospholipids (plipids) and cell wall components, considering cell wall components as the sum of teichoic acid, peptidoglycans and carbohydrates composition. The rest of components - DNA, RNA and traces-were kept fixed. The composition of proteins and plipids were randomly selected within +/-10% their current value. In this way, the total cell wall components composition was calculated following equation 2.

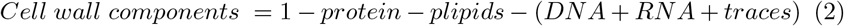

Consecutively, the value of each cell component, was distributed within teichoic acid, peptidoglycans and carbohydrates following the same proportion as they had in the original biomass synthesis reaction. For each randomly selected value, a new biomass synthesis reaction was obtained. This new biomass synthesis reaction was maximised as the objective function using FBA in COBRApy [54] maintaining fixed ethanol and CO_2_ uptake rates. We repeated this process 1000 times, so that we obtained 1000 different biomass synthesis reactions. The composition of the cell wall components, proteins and phospholipids was stored for each biomass synthesis reaction, together with the growth rate, and acetate and propionate production rate. The obtained growth rate, acetate and propionate production rate were normalised with respect the original values and were plotted against each biomass building block (Additional file 1, Fig. S1)

### Qualitative analysis of the GEM

Model simulations were done using COBRApy, version 0.15.4 [54], IBM ILOG CPLEX 128 and Python 3.6.9. Growth capabilities were assessed using FBA. The uptake rate was constrained to be 30 mmol gDW^-1^ h^-1^ for single carbon sources and a total of 30 mmol gDW^-1^ h^-1^ for double carbon sources, unless specified otherwise. The biomass synthesis reaction was used as the objective function. Growth was considered when growth rate was larger than 0.0001 h^-1^. To calculate the product profile, the fluxes compatible with the applied constraints were sampled using the *sample* function with the ‘achr’ method in the flux_analysis submodule of COBRApy [55]. The lower bound of the biomass synthesis reaction was constrained to be at least 99% of the maximum growth rate calculated by FBA. Presented results are the average and standard deviation based on 5000 iterations generated at each condition.

To assess the fermentation of ethanol and acetate in steady-state, we forced the reaction ‘EX_cpd00029_e0’ to uptake acetate instead of producing it as it was originally defined in the model. Therefore we added three reactions and one metabolite in the model to account for the production of acetate. The added metabolite is ‘totalacetate_t0’ and represents total acetate. The first reaction represents transport of extracellular acetate to total acetate (trans_ac_e0: ac_e0 → totalacetate_t0). The second reaction represents transport of intracellular acetate to total acetate (trans_ac_c0: ac_c0 → totalacetate_t0). The third reaction represents the production and consumption of intracellular and extracellular acetate, which results in total acetate (EX_AC_t0: totalacetate_t0 ↔ ac_e0 + ac_c0).

### Quantitative analysis of the GEM

The reconstructed GEM iANEO_SB607 was subjected to dFBA to simulate batch growth of *A. neopropionicum* on ethanol and ethanol plus acetate. In contrast to the qualitative study, no additional reactions were added to simulate batch growth on ethanol plus acetate. Therefore, the extracellular reaction of acetate (EX_cpd00029_e0) was unconstrained to allow consumption or production of acetate. To constrain the feasible flux space, ethanol uptake was specified to follow a Michaelis-Menten-like kinetics (Eq. 3) with parameters q_Si,max_ and K_m,i_:

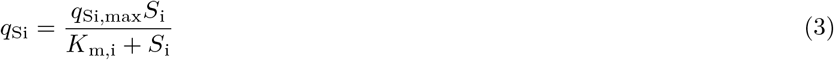

where q_Si_ is the uptake rate of substrate *i* (mmol gDW^−1^ h^−1^); q_Si,max_ is the maximum uptake rate of substrate *i* (mmol gDW^−1^ h^−1^); K_m,i_ is the Michaelis-Menten constant (mM) for substrate *i* and S_i_ is the concentration of substrate *i* (mM). K_m,i_ was determined based on experimental data and model fitting (Additional file 1, Table S1). q_Si,max_ was calculated from experimental data of batch fermentations. Concentrations of substrates, products and biomass over time were determined as follows. First, the Vs_i_ was calculated using Eq. (3) for each given time step and the defined initial concentrations. Then, FBA was applied under those constraints to compute the fluxes at maximum growth rate. After that, the following ordinary differential equations (ODE) were solved:

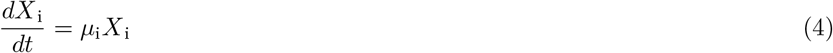

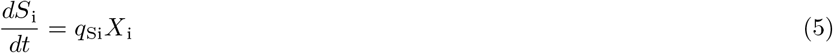

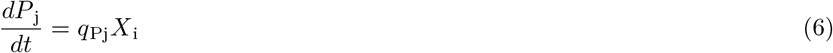

where X_i_ is the biomass concentration (g L^-1^); *µ* is the specific growth rate (h^-1^); S_i_ is the concentration of substrate *i* (mM); q_Si_ is the uptake rate of substrate *i* (mmol gDW^−1^ h^−1^); q_Pj_ is the production rate of product *j* (mmol gDW^−1^ h^−1^), and P_j_ is the concentration of product *j* (mM). Equations 4,5 and 6 were used to calculate X, S_i_ and P_j_. S_i_ is used as input to calculate the next state following equation 3. For each time step, the concentration of biomass, substrate and products were stored and plotted.

### Experimental batch fermentation data

#### Cultivation conditions

*A. neopropionicum* DSM 3847^T^ was obtained from the German Collection of Microorganisms and Cell Cultures (DSMZ, Braunschweig, Germay). Batch fermentations were done in 117 mL serum bottles containing 50 mL medium with the following composition (per litre): 0.9 g NH_4_Cl, 0.3 g NaCl, 0.8 g KCl, 0.2 g KH_2_PO_4_, 0.4 g K_2_HPO_4_, 0.2 MgSO_4_ x 7 H_2_O, 0.04 CaCl_2_ x 2 H_2_O, 3.36 g NaHCO_3_, 10 mL trace element solution from DSM medium 318, 1 mL vitamin solution, 0.5 g yeast extract, 0.3 g Na_2_S x *x* H_2_O (*x*=9-11) as reducing agent and 0.5 mg resazurin as redox indicator. The vitamin solution contained (per liter): 0.5 g pyridoxine, 0.2 g thiamine, 0.2 g nicotinic acid, 0.1 g p-aminobenzoate, 0.1 g riboflavin, 0.1 g pantothenic acid, 0.1 g cobalamin, 0.05 g folic acid, 0.05 g thioctic acid and 0.02 g biotin. The headspace of the bottles was filled with a gas mixture of N_2_/CO_2_ (80:20 % v/v; 170 kPa). To test growth in the presence of H_2_, the headspace of bottles was filled instead with a gas mixture of H_2_/CO_2_/N_2_ (10:20:70 and 80:20:0 % v/v; 170 kPa). Growth was assessed on the following substrates: ethanol, lactate, glucose and xylose, at an initial concentration of 25 mM. Where indicated, acetate (10 and 25 mM) was added to ethanol-fed cultures. The pH of the medium was 7.1 - 7.2. Cultures were incubated at 30°C statically.

#### Analytical techniques

Liquid and headspace samples were taken periodically over the course of batch fermentations and analysed for biomass, substrate and product concentrations. Biomass growth was measured by optical density at 600 nm (OD_600_). Biomass concentration (mg_CDW_ L^-1^) was estimated from OD_600_ measurements using the correlation: mg_CDW_ L^-1^ = (OD_600_ - 0.016)/0.0032, which was experimentally determined from *A. neopropionicum* cultures grown on ethanol. Concentrations of soluble compounds in the supernatant of liquid samples were determined using high-pressure liquid chromatography (HPLC) (LC-2030C Plus, Shimadzu, USA). The HPLC was equipped with a Shodex SH1821 column operated at 65°C. A solution of 0.1 N H_2_SO_4_ was used as mobile phase, at a flowrate of 1 mL/min. Detection was done via a refractive index detector. Concentrations below 0.2 mM could not be accurately quantified and are considered traces. Concentrations of gases in headspace samples were determined via gas chromatography (GC) (Compact GC 4.0, Global Analyser Solutions, The Netherlands). To analyse H_2_, a Molsieve 5A column operated at 140°C coupled to a Carboxen 1010 column was used. CO_2_ was analysed in a RT-Q-BOND column at 60°C.

## Results

### Reconstruction of iANEO_SB607, the first GEM of *A. neopropionicum*

A draft model of the metabolism of *A. neopropionicum* was developed by automatic reconstruction using the publicly available genome sequence of the microorganism (DDBJ/EMBL/GenBank accession number: LRVM00000000; [31]). The draft model comprised 491 genes, 855 metabolites and 907 reactions. This preliminary model predicted growth only on rich medium supplemented with amino acids and biomass precursors, and it did not predict the production of propionate and acetate. We performed an extensive manual curation process that resulted in the deletion, modification or addition of reactions, metabolites and genes (see git repository). The final model, iANEO_SB607, comprises 607 genes, 815 metabolites and 932 reactions (Table 1). This is the first GEM of the propionigenic bacterium *A. neopropionicum*.

**Table 1.** Composition of iANEO_SB607.

| Features | Amount |
| --- | --- |
| **Genes** | **607** |
| **Metabolites** | **815** |
| Intracellular metabolites | 742 |
| Extracellular metabolites | 73 |
| **Reactions** | **932** |
| Metabolic reactions | 771 |
| Transport reactions | 88 |
| Exchange reactions | 73 |
| Reactions associated with genes | 733 |
| Reactions non-associated with genes | 199 |

Two compartments are recognised in the model: the intracellular compartment (id: ‘c0’) and the extracellular compartment (id: ‘e0’). Metabolites are assigned to either one of the compartments. Reactions are classified as metabolic reactions, transport reactions and exchange reactions. Metabolic reactions describe the biochemical conversion of metabolites within the intracellular compartment. Transport reactions describe the transport of metabolites across the intracellular and extra-cellular compartments. Exchange reactions simulate the excretion of metabolites outside the cell or the uptake of metabolites into the cell. Reactions are distributed within cell subsystems (Fig. 1). More than 60 % of reactions belong to the central carbon metabolism (128), amino acid metabolism (148), biosynthesis of biomass building blocks (214) and transport reactions (88). The model also includes reactions involved in the production of acetate, propionate, butyrate, propanol, isobutyrate and isovalerate. Approximately 80 % of reactions could be associated to genes present in the genome of *A. neopropionicum*. The remaining 20 % of reactions are not associated with genes; these are mostly exchange reactions, diffusion transport reactions, spontaneous reactions or general reactions describing, in a summarised manner, the biosynthesis of biomass building blocks (e.g., lipids, carbohydrates).

**Figure 1.**
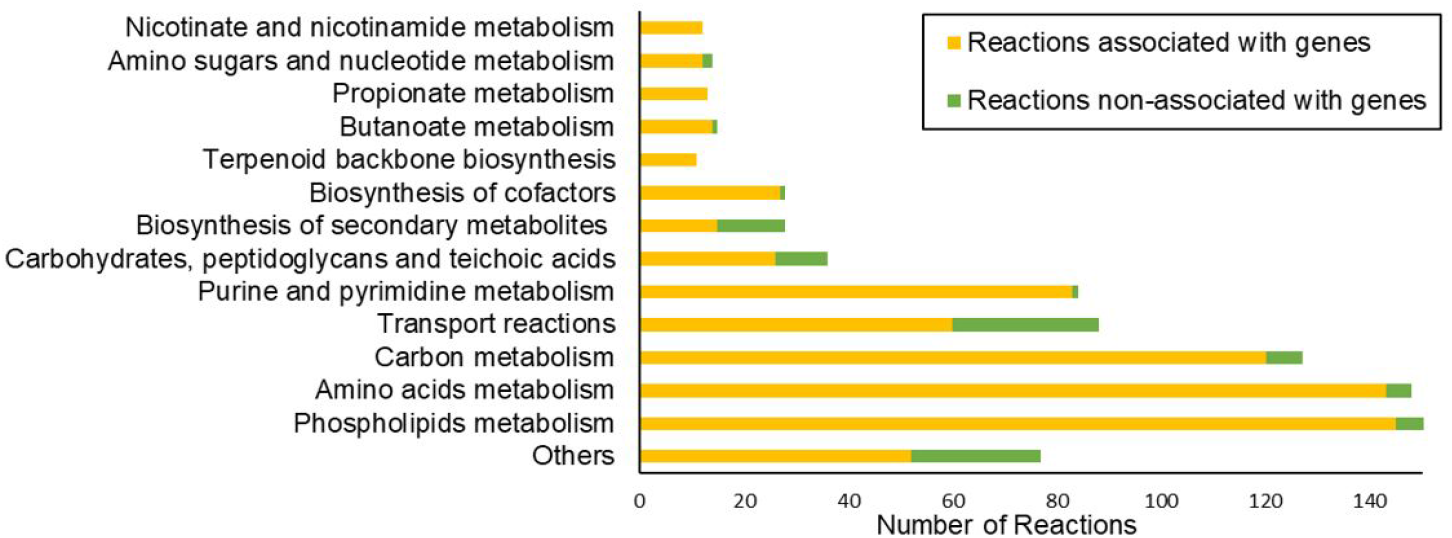
Distribution of the reactions of the iANEO_SB607 model within cellular subsystems

### Sensitivity analysis of the biomass synthesis reaction

The constructed biomass synthesis reaction (BIOMASS_Aneopro_w_GAM) accounts for the production of DNA, RNA, proteins, peptidoglycans, phospholipids, teichoic acids and the solute pool. It also includes the growth-associated ATP maintenance (GAM) as an hydrolysis reaction, and the non-growth associated ATP maintenance (NGAM) as a reaction of ATP phosphohydrolase (rxn00062_c0). The lower bound of this reaction was constrained to a rate of 8.4 mmol ATP g_DW_^-1^ h^-1^, an estimation based on the models of *C. beijerinckii* [51] and *C. autoethanogenum* [52].

Since the biomass synthesis reaction of *A. neopropionicum* was developed based on these two other species, we performed a sensitivity analysis to test its robustness. The analysis showed the effect of modifying the proportion of the main biomass components from the biomass synthesis reaction on model predictions (i.e., growth and production rates). In all scenarios tested, growth and production rates remained virtually unaffected (Additional file 1, Fig. S1). The largest deviation of the growth rate, acetate and propionate production rates were *±* 0.0005 h^-1^, *±* 0.005 mmol g_DW_^-1^ h^-1^ and *±* 0.0025 mmol g_DW_^-1^ h^-1^, respectively, which are negligible. The biomass synthesis reaction was therefore considered a reliable representation of the biomass composition of *A. neopropionicum*.

### Quality of the GEM iANEO_SB607

The quality of the iANEO_SB607 model was evaluated using the SBML validator [56] and the test suite Memote [57]. The GEM was correctly defined in SBML format, level 3, version 1. The GEM obtained an overall Memote score of 72 %. All metabolites, reactions and genes were fully annotated. The annotation per database of reactions and metabolites scored 83 %, however the annotation per database of genes scored a much lower value, 33 %. Reactions are mass and charge balanced, except for reactions associated to the synthesis of biomass precursors. The model does not have infeasible cycles and all metabolites are connected. However, the model is only partly stoichiometrically consistent (55 % scoring); this is due to the creation of metabolites to account for biomass precursors. These metabolites (e.g., RNA) lack a defined formula or a correct charge and, thus, their associated reactions are considered stoichiometrically inconsistent.

### Qualitative assessment of iANEO_SB607 through analysis of growth phenotypes

The iANEO_SB607 model was qualitatively validated by assessing growth of *A. neopropionicum* on several carbon sources and contrasting the results with experimental data. Model predictions matched most of the growth phenotypes observed in cultivation experiments from this and previous studies (Table 2; full data is available in the git repository and Additional file 1, Table S2).

**Table 2.** Growth phenotypes of *A. neopropionicum* on different substrates, predicted by the iANEO_SB607 model and observed in experiments from this and previous studies. +, Positive; -, negative; w, weakly positive; ND, no available data.

| Substrates | iANEO_SB607 | This study (exp.) | [17] | [9] | [15] |
| --- | --- | --- | --- | --- | --- |
| Ethanol | + | + | + | + | + |
| Ethanol & Acetate | + | + | + | ND | ND |
| Ethanol & Alanine | +<sup>[a]</sup> | ND | ND | ND | + |
| Ethanol & Serine | +<sup>[b]</sup> | ND | ND | ND | + |
| Pyruvate | + | ND | + | + | ND |
| D-Lactate | + | + | + | w<sup>[c]</sup> | ND |
| D-Glucose | + | + | + | - | ND |
| Xylose | + | + | + | + | ND |
| L-Threonine | + | ND | + | + | ND |
| L-serine | + | ND | + | + | ND |
| L-Alanine | + | ND | + | + | ND |
| D-Alanine | + | ND | + | ND | ND |
| L-Valine | - | ND | - | w | ND |
| L-Leucine | - | ND | ND | w | ND |
| L-Isoleucine | - | ND | ND | w | ND |
| Lysine | - | ND | - | ND | ND |
| L-Proline | - | ND | - | - | ND |

<sup>[a]</sup> L-Alanine
<sup>[b]</sup> L-Serine
<sup>[c]</sup> (L-D)-Lactate

The model predicts growth of *A. neopropionicum* on ethanol. Growth on xylose and on glucose is also predicted by the model and supported by experimental evidence, with exception of one study, which reported no growth of *A. neopropionicum* on glucose [9]. According to a previous work, *A. neopropionicum* can also grow on D-lactate, but not on L-lactate [17]. In our batch cultivations with DL-lactate as substrate, we repeatedly observed that only ≈ 50 % of the substrate was used. The purity of the L- enantiomer in the racemic mixture solution was, according to the manufacturer, 27 - 33 %. This indicates that D-lactate is indeed used by *A. neopropionicum* but it does not exclude the possibility that L-lactate is also metabolised. Yet, since the latter could not be confirmed, the model considers only the utilisation of D-lactate. The model predicts growth on pyruvate as well as on one pyruvate-derived amino acid, alanine. Serine also supports growth of *A. neopropionicum*, as predicted by the model and observed in cultivation experiments. The model indicates that branched-chain amino acids (valine, leucine and isoleucine) as well as TCA-derived amino acids (lysine and proline), with exception of threonine, are not utilised.

Further model validation was performed by assessing the product profile on a number of substrates from which sufficient experimental data was available, specifically: ethanol, lactate, glucose, xylose, L-threonine, L-serine, L-alanine, ethanol plus acetate, ethanol plus L-serine and ethanol plus L-alanine. The model predicted propionate and acetate as the major fermentation end products with all the substrates tested, in accordance with experimental evidence (Fig. 2; full data is available in the git repository and Additional file 1, Table S2).

**Figure 2.**
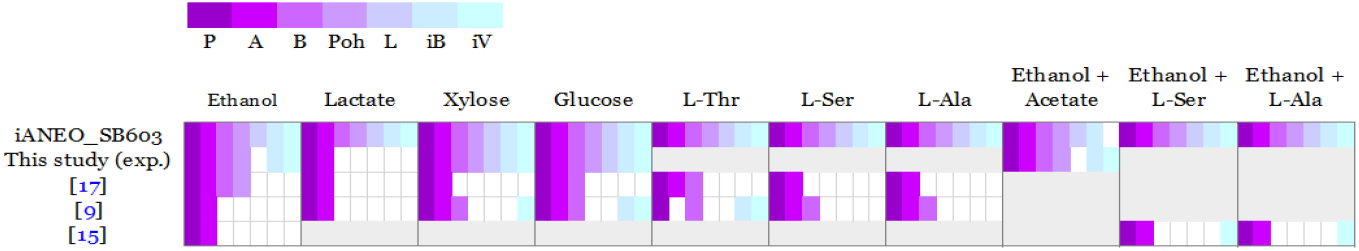
Product profile of the fermentation of different substrates by *A. neopropionicum*, predicted by the GEM iANEO_SB607 and observed in experiments from this and previous studies. P: Propionate; A: acetate; B: butyrate; Poh: propanol; L: lactate; iB: isobutyrate and iV: isovalerate. White spaces indicate the product is not reported produced. Grey areas indicate no available data.

Butyrate, propanol, lactate, isobutyrate and isovalerate are also predicted by the model as fermentation products in all cases, albeit in different proportions. Butyrate appears as a minor product in all the simulations and cultivation experiments, except for in the fermentation of L-threonine; in this case, the model predicts butyrate as a major end product, as previously reported [9]. According to model simulations and in agreement with our experimental data, lactate, an intermediate of the acrylate pathway, and propanol are produced in minor amounts. In batch cultivations carried out in this study, isobutyrate and isovalerate were detected as traces with ethanol (plus acetate), glucose or xylose as substrates, but not with lactate. The model predicted both products to be produced as traces with these substrates. Model simulations predicted enhanced production of isobutyrate and isovalerate with ethanol plus L-valine and ethanol plus L-leucine as substrates, respectively (not shown), as observed in one study [9]. The model also predicted the production of isovalerate when L-alanine or L-serine are co-substrates with ethanol, which is in agreement with observations from a recent work [15].

H_2_ was not detected as product in any of the fermentations of *A. neopropionicum* carried out in this study (with substrates: ethanol (plus acetate), lactate, glucose, xylose). In addition, H_2_ was not utilised nor affected the growth or the product profile of *A. neopropionicum* cultures growing on ethanol (Additional file 1, Fig. S2). Previous works reported the same observations [17, 58]. A ferredoxin hydrogenase is annotated in the genome of *A. neopropionicum* (CLNEO_18070; EC 1.12.7.2; model id:’rxn05759_c0’); yet, given the collected evidence, this reaction was blocked in the model.

### Quantitative assessment of iANEO_SB607 through dFBA

The iANEO_SB607 model of *A. neopropionicum* was evaluated quantitatively by simulating the dynamics of batch fermentation using dFBA. Three conditions were considered, with regard to the substrates present: 25 mM ethanol, 25 mM ethanol plus 10 mM acetate, and 25 mM ethanol plus 25 mM acetate. To constrain the model, we used empirical data of ethanol consumption, product formation and cell growth from cultivation experiments. The fermentation profiles obtained by dFBA were contrasted with the experimental data of batch incubations.

For the condition with only ethanol (and CO_2_) as substrate, the time-course data obtained through dFBA accurately reproduced the fermentation profile, with only small deviations (Fig. 3). Exponential growth of *A. neopropionicum* began after a relatively short lag phase of ≈ 13 hours. During the exponential phase, ethanol was uptaken (together with CO_2_; not shown) at an empirical maximum consumption rate (q_S,max_) of 36.2 *±* 5.5 mmol ethanol g_DW_^-1^ h^-1^. Modeled ethanol consumption fitted the experimental data with a small margin of error. Propionate and acetate were produced simultaneously during the exponential phase, at empirical maximum production rates (q_P,max_ and q_A,max_) of 12.0 *±* 0.1 mmol propionate g_DW_^-1^ h^-1^ and 8.6 *±* 0.5 mmol acetate g_DW_^-1^ h^-1^, respectively. The production profile of propionate was well predicted by dFBA, estimating a final propionate concentration (10.9 mM) close to the experimental value (9.5 mM). However, dFBA predicted a final concentration of acetate (11.5 mM) moderately higher than experimentally observed (8.6 mM). The empirical maximum specific growth rate of *A. neopropionicum* (*µ*_max_) was 0.082 *±* 0.006 h^-1^ (duplication time = 8.4 h), which was used to constrain the model. In incubations, the biomass concentration peaked (44.7 *±* 1.3 mg_DW_ L^-1^) at ≈ 47 hours, and decreased afterwards. The simulation predicted a slightly deviated pattern of biomass formation during the exponential phase, and it did not predict the observed drop in the stationary phase. Yet, the predicted maximum biomass concentration (44 mg_DW_ L^-1^) matched the empirical value. Propanol (1.3 mM) and butyrate (1 mM) were detected as minor products in batch incubations; the evolution of both products was predicted correctly by the dFBA simulations. Traces of isobutyrate and isovalerate were also detected and predicted by dFBA (not shown).

**Figure 3.**
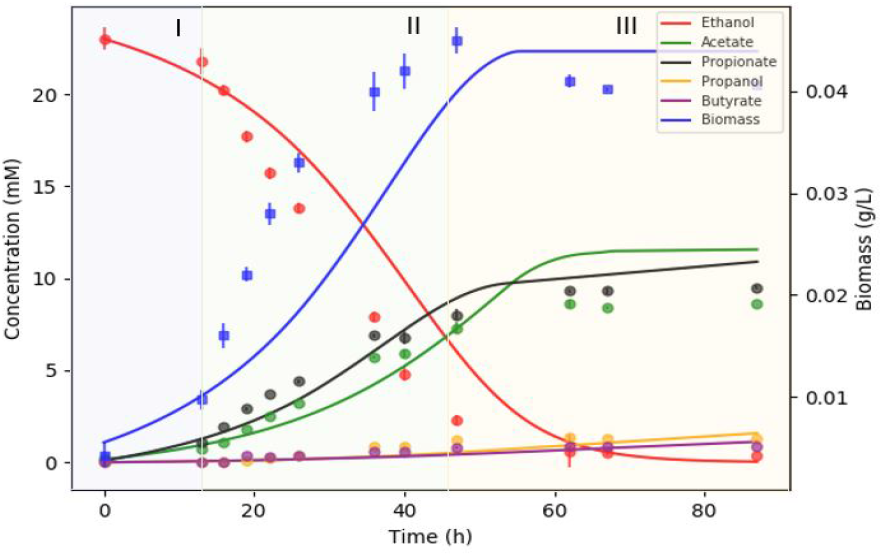
Fermentation of ethanol (25 mM) by *A. neopropionicum* in batch cultivation. Dots indicate experimental data and solid lines indicate the result of dFBA. Background colours distinguish fermentation phases: lag (blue), exponential (green) and stationary (orange).

To further evaluate the ability of *A. neopropionicum* to upgrade dilute ethanol streams from syngas fermentation, we considered a scenario with ethanol and acetate as co-substrates. Acetate is produced by acetogens as a major product of autotrophic metabolism, and it is therefore found in variable proportions in syngas fermentation effluent. To investigate the effect of acetate as co-substrate on ethanol-fermenting cultures of *A. neopropionicum*, incubations were set up with ethanol (25 mM) and acetate (10 and 25 mM) as susbtrates, and dFBA was used to simulate the dynamics of these fermentations. dFBA reproduced with high accuracy the fermentation profile of incubations containing ethanol plus 10 mM acetate (Fig. 4). In this condition, the observed *µ*_max_ was 0.098 *±* 0.005 h^-1^ (duplication time = 7.1 h); 19 % higher than in the incubations without acetate. However, less biomass was formed in comparison; the maximum biomass concentration was 41.1 *±* 0.8 mg_DW_ L^-1^ (≈ 9% lower), which was also predicted by dFBA. The presence of 10 mM acetate also affected the consumption and production rates; ethanol consumption was faster than in the absence of acetate; the q_S,max_ was 43.3 *±* 4.3 mmol ethanol g_DW_ h^-1^, a 20 % increase. The q_A,max_ in this condition dropped to 3.1 *±* 0.6 mmol acetate g_DW_ h^-1^. The biggest difference was in the q_P,max_, which was 16.4 *±* 0.8 mmol propionate g_DW_ h^-1^, a 37 % increase compared to the condition without acetate. The final propionate concentration was also slightly higher, 11.3 mM (vs. 9.5 mM). Here, again, the simulation predicted a similar propionate concentration to the observed value (12.2 mM), and a higher final acetate concentration (18.3 mM) than observed (16.7 mM). The incubations containing 25 mM acetate at the start followed a different trend than the incubations with 10 mM acetate (fermentation profile not shown). In batch bottles, the biomass concentration reached a similar value to obtained in the condition with 10 mM acetate, but the *µ*_max_, q_P,max_ and q_A,max_ were similar to the condition without acetate (data not shown). The final propionate concentration was 12.5 mM, the highest of the three conditions tested.

**Figure 4.**
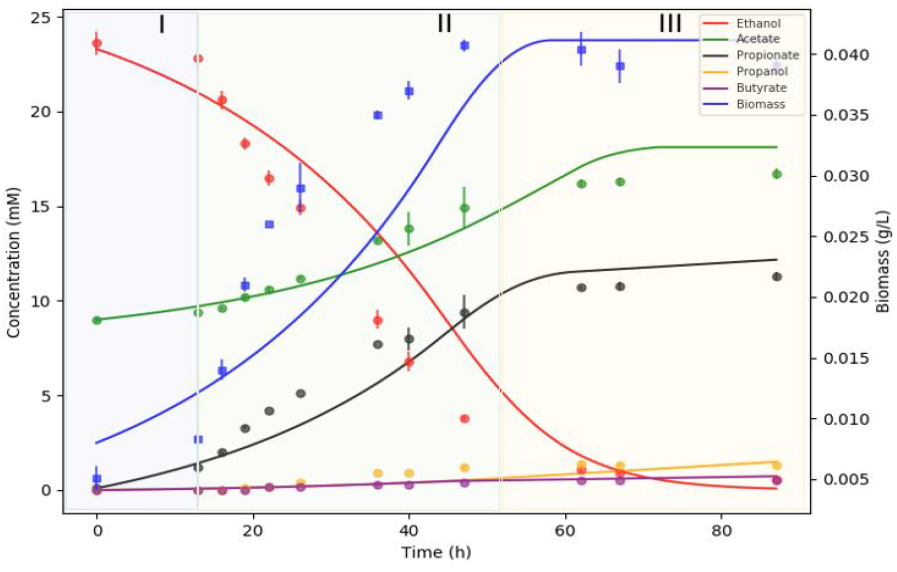
Fermentation of ethanol (25 mM) and acetate (10 mM) by *A. neopropionicum* in batch cultivation. Dots indicate experimental data and solid lines indicate the result of dFBA. Background colours distinguish fermentation phases: lag (blue), exponential (green) and stationary (orange).

The presence of acetate had an effect on the utilisation of ethanol by *A. neopropionicum*, which is reflected in the fermentation yields. The biomass yield (Y_X/S_) was slightly lower in the presence of both 10 and 25 mM acetate (1.4 g_DW_ mol ethanol^-1^ vs. 1.6 g_DW_ mol ethanol^-1^ when no acetate was present). With acetate present at the start of incubations, more ethanol was invested in propionate production, as indicated by the propionate yields (Y_P/S_, mol mol^-1^), which were 0.33, 0.38 and 0.42 for the conditions with no acetate, 10 mM acetate and 25 mM acetate, respectively. The production of acetate followed the inverse trend; acetate yields (Y_A/S_, mol mol^-1^) were 0.29, 0.18 and 0.06 for the conditions with no acetate, 10 mM acetate and 25 mM acetate, respectively. Similarly, lower yields were obtained for propanol and butyrate when acetate was present (data now shown).

### Ethanol fermentation via the acrylate pathway

The reconstructed iANEO_SB607 model describes the metabolism of ethanol fermentation and propionate production via the acrylate pathway in *A. neopropionicum* (Fig. 5). Model simulations provided new insights into the enzymatic reactions involved in propionate formation, cofactor regeneration and the energy metabolism of the cell.

**Figure 5.**
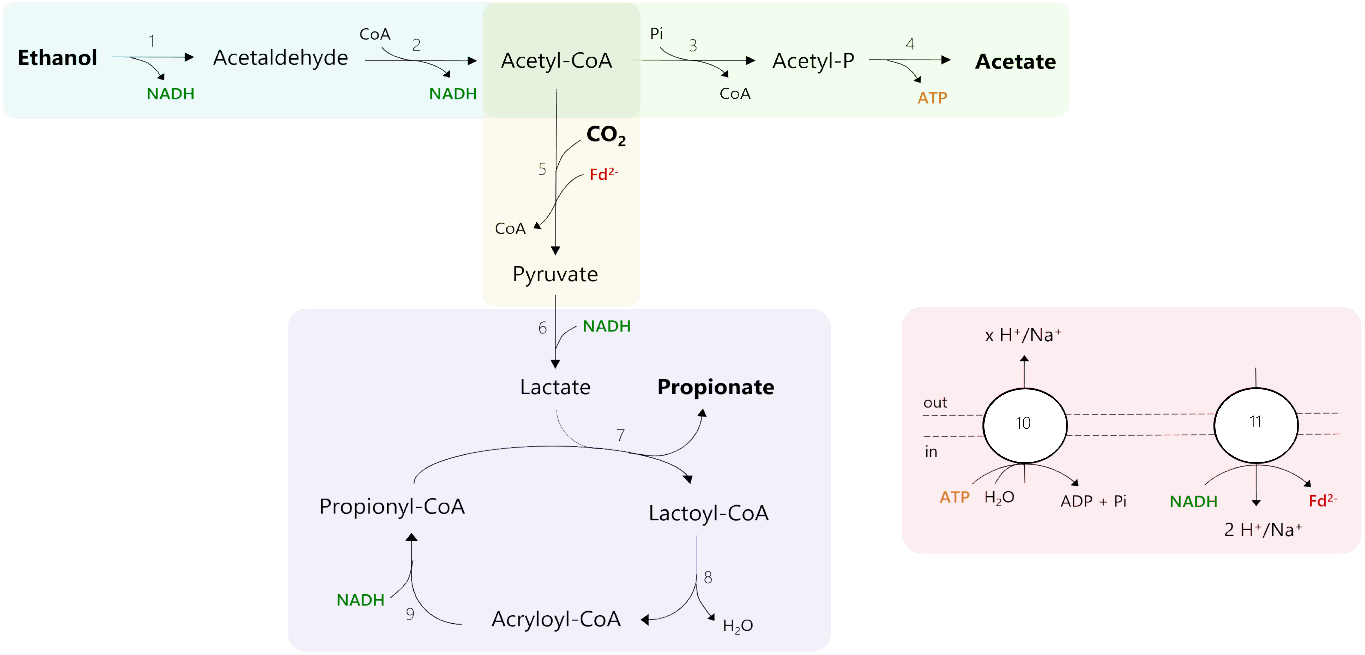
Proposed metabolism of ethanol fermentation to propionate via the acrylate pathway in *A. neopropionicum*. Coloured areas designate the following modules: ethanol oxidation (blue), acetate production (green), pyruvate synthesis (yellow), lactate production and acrylate pathway (purple), redox cofactor regeneration and ATPase (red). Numbers in reactions correspond to the following enzymes and reaction ids in the model: 1, alcohol dehydrogenase (rxn00543_c0); 2, alcohol-acetaldehyde dehydrogenase (rxn00171_c0); 3, phosphate acetyltransferase (rxn00173_c0); 4, acetate kinase (rxn00225_c0); 5, pyruvate:ferredoxin oxidoreductase (PFOR; rxn05938_c0); 6, NAD-dependent D-lactate dehydrogenase (rxn00500_c0); 7, propionate-CoA:lactoyl-CoA transferase (rxn01056_c0); 8, lactoyl-CoA dehydratase (rxn02123_c0); 9, acryloyl-CoA reductase (rxn40050_c0); 10, ATPase (rxn10042_c0); 11, Rnf complex (Rnf_c0).

Ethanol is oxidised to acetyl-CoA via acetaldehyde through alcohol and acetaldehyde dehydrogenases. The genome of *A. neopropionicum* harbours a bifunctional NAD^+^-dependent alcohol-aldehyde dehydrogenase (AdhE; CLNEO_13930) that can catalyse this two-step conversion. According to our model, two other alcohol dehydrogenases, encoded by *adh* (CLNEO_16910) and *adhB* (CLNEO_00480), could also drive the oxidation of ethanol to acetaldehyde. Initially, the model also predicted this reaction to be catalysed by NAD(P)H-dependent butanol dehydrogenase (BdhA), encoded by *bdhA* (CLNEO_09740; rxn00536_c0). However, the well-characterised BdhA of *Clostridium acetobutylicum*, which shares 60.7 % identity with that of *A. neopropionicum*, is known to contribute primarily to butanol production and it is the alcohol dehydrogenase least involved in ethanol metabolism [59]. Thus, we reasoned that BdhA would likely not be involved in ethanol oxidation in *A. neopropionicum* and excluded this reaction from model simulations.

Acetyl-CoA is partly used in the reductive reactions of the metabolism and partly invested in the formation of acetate, an energy-generating step. Acetate is synthesised via phosphate acetyltransferase (Pta; CLNEO_28570) and acetate kinase (Ack; CLNEO_28580), yielding ATP via substrate-level phosphorylation (SLP). In the reductive path, acetyl-CoA is converted to pyruvate through the CO_2_-fixating reaction catalysed by pyruvate:ferredoxin oxidoreductase (PFOR; CLNEO_15240 or CLNEO_19010 or CLNEO_17780 or CLNEO_03040 or CLNEO_04330 or CLNEO_24550). This conversion requires reduced ferredoxin (Fd^2-^) as electron carrier. Our hypothesis, supported by model predictions, is that Fd^2-^ is produced in the Na^+^-translocating ferredoxin:NAD^+^ oxidoreductase (Rnf) complex. The Rnf complex is a membrane-bound respiratory enzyme involved in energy conservation in anaerobic microorganisms [60]. During growth on high-energy substrates, it catalyses the exergonic reduction of NAD^+^ with electrons from Fd^2-^ coupled to the translocation of two cations (H^+^ or Na^+^) across the membrane. The electrochemical potential established by the Rnf complex can then be used by a membrane-bound ATP synthase for energy generation. The Rnf complex can also operate in the reverse direction to produce Fd^2-^ at the expense of ATP [61].

The genome of *A. neopropionicum* harbours a complete *rnf* cluster, composed of the genes *rnfA* (CLNEO_01390), *rnfB* (CLNEO_01400), *rnfC* (CLNEO_01350), *rnfD* (CLNEO_01360), *rnfE* (CLNEO_01380) and *rnfG* (CLNEO_01370). With ethanol as substrate, our assumption is that the Rnf complex of *A. neopropionicum* operates in reverse, generating Fd^2-^. The endergonic reduction of ferredoxin (E_o_’= - 500 to - 420 mV) with NADH (E_o_’= - 320 mV) is driven by reverse electron transport across the membrane which, in turn, is an energy-driven process. A membrane-bound V-type ATPase is present in the genome of *A. neopropionicum*, encoded by the genes *atpA/ntpA* (CLNEO_280), *atpB/ntpB* (CLNEO_290), *ntpC*, (CLNEO_260), *atpD/ntpD* (CLNEO_23400), *atpE* (CLNEO_250), *ntpG* (CLNEO_270), *ntpK* (CLNEO_240) and *ntpI* (CLNEO_23330). We theorise that ATP is hydrolysed in the ATPase to create a proton- or sodium-motive-force that is used by the Rnf complex to catalyse the reduction of ferredoxin. The production of Fd^2-^ is an energy costly process, the implications of which are addressed later in this section.

Pyruvate produced by the PFOR is subsequently reduced to lactate with NADH via D-lactate dehydrogenase (CLNEO_28010). We assumed NADPH is not used as electron carrier in this reaction, since lactate dehydrogenases have a strict specificity for NAD^+^/NADH [62, 63]. Lactate then enters the acrylate pathway, a cyclic chain of reactions involving the intermediates lactoyl-CoA, acryloyl-CoA and propionyl-CoA. The characteristic enzyme of this pathway is propionate-CoA:lactoyl-CoA transferase (Pct, EC 2.8.3.1), which exchanges the CoA moiety between propionyl-CoA and lactate, generating lactoyl-CoA and propionate as end product [64, 65]. Our first annotation of the genome of *A. neopropionicum* did not include Pct. However, an acetate CoA-transferase was present, encoded by the gene *ydiF* (CLNEO_17700), that shared 96 % identity with the purified and well characterised Pct of *A. propionicum* [65]. Thus, we deduced that *ydiF* encodes for Pct in *A. neopropionicum* and included this reaction in the model. Lactoyl-CoA dehydratase (CLNEO_17710 and CLNEO_17720) catalyses the dehydration of lactoyl-CoA to acryloyl-CoA, which is subsequently reduced to propionyl-CoA by acryloyl-CoA reductase. Our genome annotation revealed that the acryloyl reductase of *A. neopropionicum* forms an enzymatic complex with an electron-transferring flavoprotein (EtfAB). The complex, hereafter named acryloyl-CoA reductase-EtfAB (Acr-EtfAB), is also present and has been well characterised in *A. propionicum* [66]. Three gene clusters predicted to encode for acryloyl-CoA reductase (*acrC*) or EtfAB (*acrA,acrB*) were found in the genome: (i) CLNEO_21740 (*acrC*), CLNEO_21750 (*acrB_1*) and CLNEO_21760 (*acrA*); (ii) CLNEO_26130 (*acdA_1*) and CLNEO_26120 (*acrB_2*); and (iii) CLNEO_29850 (*acdA_2*) and CLNEO_29840 (*acrB_3*). The *acdA_1* and *acdA_2* genes encode for acyl-CoA dehydrogenases that share low identity (46 and 54 %, respectively) with the acryloyl-CoA reductase encoded by *acrC*; thus, we assumed that the former two enzymes are not responsible for acryloyl-CoA reductase activity. The first cluster is the only complete one, composed of acryloyl-CoA reductase (*acrC*) and the A (*acrA*) and B (*acrB_1*) subunits of EtfAB. The proteins encoded by these three genes share an identity of 92.9 %, 89.7 % and 89.1 %, respectively, with their homologues from the Acr-EtfAB complex of *A. propionicum*. The Acr-EtfAB of *A. propionicum* is a non-bifurcating soluble enzyme that catalyses the irreversible reduction of acryloyl-CoA to propionyl-CoA with NADH via electron transfer to a flavin moiety and appears not to be involved in energy conservation [66, 67]. Given their high similarity, we deduced the same features apply to the Acr-EtfAB of *A. neopropionicum*. To our knowledge, this is the first time that the Acr-EtfAB complex is identified in this microorganism.

According to the theoretical stoichiometry, the fermentation of ethanol yields propionate and acetate in a 2:1 ratio (Eq. 1). However, this ratio is not observed in cultures of *A. neopropionicum*; rather, ethanol fermentation resulted in a ≈ 1.2:1 propionate to acetate ratio (Fig. 3 and Additional file 1, Table S2). We reasoned that the theoretical ratio cannot be achieved in *A. neopropionicum* due to energetic constraints of the cell, specifically, due to the requirement of Fd^2-^. Model simulations were performed to confirm this. The oxidation of three moles of ethanol generates six moles of NADH and three moles of acetyl-CoA. To fit the theoretical 2:1 propionate to acetate ratio, two moles of acetyl-CoA would have to be used in the reductive part of the metabolism, and one mole of acetyl-CoA should be invested in acetate, with the concomitant production of one mole of ATP (via SLP). The synthesis of two moles of pyruvate from acetyl-CoA would require two moles of Fd^2-^, which is produced at the RnF complex at the expense of ATP. However, the hydrolysis of one mole of ATP (Δ*G*^*o*^ = -32 KJ mol^-1^; [68]) could drive the reduction with NADH of no more than ≈ 1.3 moles of ferredoxin (Δ*G*^*o*^ = -25 KJ mol^-1^; [69]). Moreover, two other issues arise: i) even if this one mole of ATP would solely be invested in the reduction of ferredoxin, this would leave no net ATP for growth, and ii) such a scenario would result in excess reducing equivalents from ethanol oxidation that could not be recycled in the production of propionate. Our model predictions confirmed this inconsistencies and are in agreement with the hypothesis that the propionate to acetate 2:1 ratio cannot be achieved in *A. neopropionicum* during the fermentation of ethanol. Instead, cells must invest more than one mole of acetyl-CoA in acetate production to obtain net ATP to support growth. This leaves less than two moles of acetyl-CoA available for propionate production and, overall, a propionate to acetate ratio lower than the theoretical 2:1. The actual propionate to acetate ratio (close to 1.2:1, based on the fermentation balance) depends on how much Fd^2-^ can be produced per hydrolysed ATP, which in turn depends not only on the Gibbs free energies of ATP hydrolysis and ferredoxin reduction with NADH under physiological conditions but also on the coupling ratio of the ATPase (number of cations translocated per ATP hydrolised). While the Rnf complex can be assumed to translocate two cations per ferredoxin reduced/oxidised, the coupling ratio of the ATPase remains unknown for *A. neopropionicum*. Our model fitted with a coupling ratio of the ATPase of 3 to 3.5 H^+^/Na^+^ translocated per ATP.

### Propanol and butyrate production pathways

*A. neopropionicum* cells produce propanol and butyrate as minor products of the fermentation of several substrates (Fig. 2). The model reconstruction allowed us to look into the pathways involved in propanol and butyrate production in *A. neopropionicum*. According to model predictions, propanol is formed from propionyl-CoA via propionaldehyde in a two-step reductive conversion catalysed by AdhE. Reduction of propionaldehyde can also be catalysed by the NAD^+^-dependent alcohol dehydrogenases encoded by *adh* (CLNEO_16910) and *adhB* (CLNEO_00480).

Four routes have been described for the production of butyrate in bacteria: the acetyl-CoA, glutarate, lysine, and 4-aminobutyrate/succinate pathways [70]. The acetyl-CoA pathway is dominant within members of the phylum *Firmicutes* [70], which *A. neopropionicum* belongs to. Most enzymes of the acetyl-CoA pathway were either found in the genome of *A. neopropionicum*, were assigned during the re-annotation or were identified through protein sequence alignment. In this pathway, acetyl-CoA is first converted to the intermediate butyryl-CoA in four steps. First, two acetyl-CoA molecules are condensed to yield acetoacetyl-CoA, a reaction catalysed by acetoacetyl-CoA thiolase (EC 2.3.1.9). A gene coding for this thiolase was absent in the genome of *A. neopropionicum*. However, since the rest of genes of the pathway were present, we added this reaction to the model during the gap-filling process. Acetoacetyl-CoA is reduced to 3-hydroxybutyryl-CoA via either NAD(P)-dependent 3-hydroxybutyryl-CoA dehydrogenase (EC 1.1.1.157) or NAD(P)-dependent acetoacetyl-CoA reductase (EC 1.1.1.36). In *A. neopropionicum*, this activity could be catalysed by an enzyme annotated as 3-oxoacyl reductase (EC 1.1.1.100; CLNEO_27400 or CLNEO_28900 or CLNEO_06910 or CLNEO_18000 or CLNEO_02960 or CLNEO_14220), which showed 54 - 66 % identity with the characterised acetoacetyl-CoA reductase of *Cupriavidus necator*. In *C. necator* and in *A. neopropionicum*, this enzyme is annotated as NADP-dependent. Next, crotonyl-CoA is formed via a dehydration reaction catalysed by 3-hydroxybutyryl-CoA dehydratase (EC 4.2.1.55). *A. neopropionicum* harbours a 3-hydroxyacyl dehydratase (EC 4.2.1.59; CLNEO_27430) that, according to database searches (BRENDA and MetaCyc), could catalyse this reaction. Crotonyl-CoA is reduced to butyryl-CoA via the butyryl-CoA dehydrogenase/electron-transferring flavoprotein complex (Bcd-EtfAB). The Bcd-EtfAB complex is an electron-bifurcating enzyme that couples the exergonic reduction of crotonyl-CoA to butyryl-CoA (E_o_’= -10 mV) by NADH to the endergonic reduction of Fd by NADH [60]. Reduced ferredoxin contributes to energy conservation by regeneration of NADH via the Rnf complex. Our model predicts that, in *A. neopropionicum*, reduction of crotonyl-CoA could be catalysed by the Acr-EtfAB complex (not involved in ferredoxin reduction) or by either of the two acyl-CoA dehydrogenases that cluster with subunits of the EtfAB complex (*acdA_1* –*acrB_2* and *acdA_2* –*acrB_3*). Among the three, the acyl-CoA dehydrogenase encoded by *acdA_2* showed the highest identity with the butyryl-CoA dehydrogenases (Bcd) of *Clostridium acetobutylicum* and of *Clostridium kluyveri* (64 and 63 %, respectively). It remains a question whether, in *A. neopropionicum*, these complexes could be involved in the reduction of ferredoxin.

Two distinct routes have been characterised for the conversion of butyryl-CoA to butyrate. The first pathway, identified in *Clostridium acetobutylicum* [71], involves phosphate butyryltransferase (Ptb; EC 2.3.1.19) and butyrate kinase (Buk; EC 2.7.2.7) and therefore yields ATP via substrate-level phosphorylation. In the second pathway, butyryl-CoA:acetate CoA-transferase (But; EC 2.8.3.8) transfers the CoA moiety of butyryl-CoA to acetate, releasing acetyl-CoA and butyrate. The co-occurrence of genes involved in both pathways is rare among butyrate producers [70]. The genome of *A. neopropionicum* does not encode for Ptb nor Buk, yet our annotation initially assigned these activities to phosphate acetyltransferase (Pta) and acetate kinase (Ack). Pta and Ack of *A. neopropionicum* are significantly similar to the well-characterised Ptb (63.8 % identity) and Buk (71 % identity), respectively, of *Clostridium acetobutylicum*. This high similarity can be expected, since these enzymes belong to the same family and have a very similar function [72]. However, it is known that carboxylate kinases differ in the size and hydrophobicity of the substrate binding site, which ultimately determines their substrate specificity [73, 74]; butyrate kinases have a wider binding site than acetate kinases to accommodate for a larger substrate. Thus, we assumed that Pta and Ack are likely not involved in butyrate production in *A. neopropionicum*, and excluded this route in model simulations. Instead, we hypothesise that butyrate production in *A. neopropionicum* takes place via butyryl-CoA:acetate CoA-transferase activity. Our model predicts that the propionate-CoA:lactoyl-CoA transferase (Pct) encoded by the gene *ydiF* catalyses this reaction. The Pct of *A. propionicum* exhibits broad substrate specificity for monocarboxylic acids, including butyrate, supporting the model prediction [65]. Also, in *Escherichia coli* K-12, the protein encoded by *ydiF* is an acyl-CoA transferase that has been shown to produce butyrate from butyryl-CoA [75]. Thus, in our model, the butyryl-CoA:acetate CoA-transferase activity is assigned to the enzyme Pct.

### Identification of the NADH-dependent reduced ferredoxin:NADP^+^ oxidoreductase (Nfn)

During the genome re-annotation and manual curation process, we identified the enzyme NADH-dependent reduced ferredoxin:NADP^+^ oxidoreductase (Nfn). Nfn is an iron-sulfur flavoprotein complex with electron-confurcating/bifurcating activity that reversibly catalyses the endergonic reduction of NADP^+^ by NADH coupled with the exergonic reduction of NADP^+^ by Fd^2-^ [76]. Nfn is composed of two subunits, NfnA and NfnB, whose coding genes were both found in the genome of *A. neopropionicum* under the locus tags CLNEO_00270 and CLNEO_00280, respectively. In the initial automatic annotation, these two genes were assigned to ferredoxin:NADP^+^ oxidoreductase and glutamate synthase, respectively. It has been reported that NfnA/B share sequence similarities with these two enzymes [76]. Upon manual inspection, we observed that the protein complex showed a significant identity (60 - 66 %) with the Nfn complexes of *Clostridium kluyveri* [77] and of *Clostridium autoethanogenum* [78], which lead us to the re-assignation of the two proteins as NfnA and NfnB.

We used modelling to look into the role of the Nfn complex in the metabolism of *A. neopropionicum* during growth on ethanol. The model shows that the Nfn generates NADPH from NADH and Fd^2-^ for NADPH-dependent reactions of the cell. For instance, NADPH is required during butyrate production in the reduction of acetoacetyl-CoA to 3-hydroxybutyryl-CoA, a reaction catalysed by a NADPH-dependent 3-oxoacyl reductase. NADPH is also required in the biosynthesis of amino acids and biomass precursors. In our model, the Nfn complex does not function in the reverse direction, the production of Fd^2-^, during growth on ethanol; this would require NADPH, and ethanol oxidation is assumed to occur only via NAD^+^-dependent reactions.

### Fermentation of other carbon sources: the case of lactate

Besides ethanol, *A. neopropionicum* can grow on lactate, sugars and some pyruvate-derived amino acids (Table 2). The fermentation of these carbon sources proceeds with key differences compared to the fermentation of ethanol. To illustrate this with an example, we chose to describe here the case of lactate fermentation, since lactate is a typical substrate of propionate-producing bacteria and, in particular, of species that use the acrylate pathway [64, 16]. The GEM iANEO_SB607 describes the pathway of lactate fermentation to propionate in *A. neopropionicum*. Lactate, when used as substrate, is consumed in both oxidative and reductive reactions. In the oxidative branch, lactate is oxidised to pyruvate via lactate dehydrogenase, generating NADH. Then, PFOR catalyses the decarboxylation of pyruvate to acetyl-CoA and CO_2_, a reaction that generates Fd^2-^. Here, PFOR functions in the opposite direction to what occurs in the condition with ethanol as substrate. The same occurs in the fermentation of sugars, pyruvate and pyruvate-derived amino acids. In other words, contrary to the fermentation of ethanol, the oxidation of these substrates generates directly Fd^2-^, which can contribute to energy conservation. Acetyl-CoA is used for acetate production via Pta and Ack, yielding ATP via SLP. In the reductive branch, lactate is converted to propionate via the reactions of the acrylate pathway. In this conversion, NADH is needed for the reduction of acryloyl-CoA to propionyl-CoA, but the NADH obtained in the oxidation of lactate is insufficient. Our model predicts that additional NADH is produced in the Rnf complex. Opposite to the scenario with ethanol as substrate, here the Rnf catalyses the exergonic reduction of NAD^+^ with electrons from Fd^2-^. This reaction is coupled to the translocation of two cations across the membrane, generating an ion-motive force that can be used by the ATPase to produce ATP. Thus, in the fermentation of carbon sources other than ethanol, ATP is generated both by SLP via acetate production and by chemiosmosis driven by the oxidation of Fd^2-^.

## Discussion

In this study we have presented iANEO_SB607, the first GEM of the propionate-producer *A. neopropionicum*. The overall Memote score of 72 % indicates the high quality of the model. The low score of gene annotation per database (33 %) was expected, since there were almost no available annotations of the genome of *A. neopropionicum* in public databases recognised by Memote. Our focus has been on gaining insight into the metabolism of ethanol fermentation to propionate, which in this bacterium occurs via the acrylate pathway.

Propionate production in *A. neopropionicum* proceeds via the acrylate pathway, which involves the key enzyme propionate-CoA:lactoyl-CoA transferase (Pct). We found that, in *A. neopropionicum*, Pct is encoded by the gene *ydiF* (CLNEO_17700). In addition, we revealed that acryloyl-CoA reductase, which catalyses the reduction of acryloyl-CoA to propionyl-CoA with NADH, forms a complex with an electron-transferring flavoprotein. This is the first time that the complex, Acr-EtfAB, is identified in this microorganism. Based on evidence from the Acr-EtfAB of *A. propionicum* [66], we hypothesise that this complex is not involved in energy conservation via ferredoxin reduction.

We have also addressed an important issue regarding the energetic metabolism of the *A. neopropionicum*. During growth on ethanol, Fd^2-^ is required to reduce acetyl-CoA to pyruvate. In the earliest description of the metabolism of *A. neopropionicum*, authors suggested that the oxidation of acetaldehyde proceeded with ferredoxin as electron carrier, thus fulfilling this demand [17]. However, at the present time it is acknowledged that aldehyde dehydrogenases are NAD(P)-dependent enzymes [79], which invalidates that theory. Theoretically, the Acr-EtfAB complex could drive the reduction of Fd (E_o_’= - 500 to - 420 mV) with NADH (E_o_’= - 320 mV) via electron bifurcation, given the high reduction potential of the acryloyl-CoA/propionyl-CoA pair (E_o_’= + 70 mV). The complex shares high protein sequence similarity with the electron-bifurcating Bcd-EtfAB complex of *C. kluyveri* [66, 80], which relies on the reduction potential of the crotonyl-CoA/butyryl-CoA pair (E_o_’= -10 mV) to catalyse the reduction of Fd with NADH. However, the purified Acr-EtfAB of *A. propionicum* appears not be involved in electron bifurcation [66], most likely to prevent transient accumulation of the very reactive intermediate acryloyl-CoA [81, 80]. Instead, our model predicted that the generation of Fd^2-^ occurs in the Rnf complex. This is a respiratory enzyme of anaerobes whose role was only discovered years after the description of *A. neopropionicum* [60]. Although in most anaerobes it operates in the direction of ferredoxin oxidation to drive ATP synthesis, reverse electron transport in the Rnf complex at the expense of ATP has also been reported to occur in, e.g., acetogens, during growth on low-energy substrates (i.e., ethanol, lactate) that yield insufficient or no Fd^2-^ [61]. The Rnf complex had been previously annotated in *A. neopropionicum* [31] and identified in its proteome [15]. It is also present in the close relative *A. propionicum* [29]. Here, through the re-annotation an a thorough manual curation process, we identified all its subunits (*rnfA-E, rnfG*) and via modelling we verified its involvement in the metabolism of the cell. During growth on ethanol, ATP is solely produced via SLP. Our model simulations showed that, in order to generate net ATP to sustain growth, *A. neopropionicum* must shift the theoretical propionate to acetate ratio of 2:1 (Eq. 1) towards more acetate and, therefore, more ATP. This was in agreement with the fermentation balance, which showed a propionate to acetate ratio of 1.2:1 (Fig. 3 and Additional file 1, Table S2), and with previous observations [9].

Our annotation of the genome of *A. neopropionicum* revealed the presence of another key enzyme of the metabolism of anaerobes: the Nfn complex. This electron-bifurcating enzyme had not been identified yet in this microorganism. We hypothesised that, during ethanol oxidation, the role of Nfn in *A. neopropionicum* is to produce NADPH for NADPH-dependent reactions of the metabolism, which our model confirmed. Further investigation is needed to define the instances in which the Nfn operates in the reverse direction, bifurcating electrons from NADPH to produce Fd^2-^ and NADH. The directionality and role of Nfn will depend on the cofactor requirements of the cell.

Butyrate and propanol are produced by *A. neopropionicum* as minor products during the fermentation of several substrates (Fig. 2), probably as a means to dispose of excess reducing equivalents generated during substrate oxidation. Our model showed that propanol is produced from propionyl-CoA with propionaldehyde as intermediate via NAD^+^-dependent reactions, as Tholozan et al. suggested [17]. The butyrate production pathway had not been described yet in this microorganism. Here, we showed that butyrate production occurs via the acetyl-CoA pathway, and very likely involves butyryl-CoA:acetate CoA-transferase activity instead of the enzymes phosphate butyryltransferase and butyrate kinase, which were not found in the genome. Further research should confirm whether, in *A. neopropionicum*, Pct (*ydiF*), key enzyme of the acrylate pathway, is indeed responsible for the conversion of butyryl-CoA to butyrate, as observed *in vitro* for the homologous CoA transferases of *A. propionicum* [65] and *E. coli* K-12 [75].

Another aspect of the metabolism that we aimed to clarify was the ability of *A. neopropionicum* to produce and consume H_2_. In our batch cultivations on different substrates, H_2_ was not produced nor consumed. Tholozan et al. also observed this, and reported no hydrogenase activity in cell-free extracts of *A. neopropionicum* [17]. However, in a recent study, traces of H_2_ were detected in cultures of *A. neopropionicum* cultivated in the absence of a carbon source [9]. Given this observation and the fact that a putative ferredoxin hydrogenase is present in our annotated genome, the ability of *A. neopropionicum* to produce or consume H_2_ under specific conditions shall not be entirely ruled out. Our results do confirm that neither the fermentation balance nor the growth of *A. neopropionicum* are affected by the presence of H_2_, at least with ethanol as substrate (Additional file 1, Figure S2), as reported earlier [17]. This is an advantageous trait when considering this strain for its application in syngas-fermenting co-cultures, since syngas contains H_2_. Interestingly, H_2_ tolerance is manifested differently in functionally-related strains. While *P. propionicus* and *D. propionicus* both use the methylmalonyl pathway to metabolise ethanol, the first is not affected by the presence of H_2_ while the second is strongly inhibited by it [14].

The GEM iANEO_SB607 accurately reproduced observed growth phenotypes on typical substrates (ethanol, sugars, lactate and amino acids), yet with a few discrepancies. Existing literature provided contradictory evidence regarding the ability of *A. neopropionicum* to grow on glucose and xylose [17, 11, 82, 9]. In our batch incubations, both sugars were utilised (Additional file 1, Table S2), and this is in agreement with model predictions. Therefore, we come to the conclusion that *A. neopropionicum* can use glucose and xylose for growth, and that the inconsistencies reported might be attributed to the different media used for incubations, as suggested by other authors [11]. Our results also indicate that only D-lactate, and not L-lactate, support growth of *A. neopropionicum*, as observed by Tholozan et. al [17]. Yet, the latter authors reported lactate dehydrogenase activity in cell-free extracts with D-, L- and DL-lactate, and hypothesised the presence of a lactate racemase [17]. This enzyme was absent in our annotated genome. However, *A. neopropionicum* has both L- and D-lactate dehydrogenases, so it cannot be excluded that L-lactate is also metabolised, perhaps at a much slower rate, as it has been suggested earlier [83]. The model predicts the full range of end products that *A. neopropionicum* can produce from different substrates, and estimates quantitative fluxes with a low detection limit. One particular advantage of this is that the model can identify products that are otherwise not detected in cultivation experiments. This is the case, for example, of isobutyrate and isovalerate, which are predicted as minor end products of ethanol, lactate, some amino acids and sugars fermentation, but this is not always reported *in vivo* (Fig. 2).

To further validate the model and test its applicability, we used dFBA to predict the dynamics of ethanol (plus acetate) fermentation by *A. neopropionicum* in batch cultivation. Overall, the dFBA simulations fitted well the experimental results (Fig. 3 and Fig. 4). In batch cultures with ethanol as substrate, the theoretical 2:1 molar ratio of propionate to acetate (Eq. 1) was not achieved; instead, this ratio was ≈ 1.2:1. As explained earlier, we presume this is because more ATP must be produced via SLP (and, therefore, more acetate) to sustain growth while investing part of it to drive the reduction of ferredoxin, required by PFOR. Another factor that could contribute to a ¡ 2:1 propionate to acetate ratio is the need to deal with acryloyl-CoA, a highly reactive compound; to prevent its accumulation, cells would favour the synthesis of acetate over propionate [64]. In addition, propanol was produced towards the end of batch fermentations, likely as an attempt of cells to halt further acidification of the medium. Since propanol is produced from propionyl-CoA, this would result in a lower flux towards propionate.

Interestingly, we observed that a low acetate concentration (*<* 25 mM) or low acetate:ethanol ratio (*<* 1) at the start boosted the growth rate, propionate production rate of *A. neopropionicum* during growth on ethanol. The acetate production rate was lower than in the condition without acetate at the start, suggesting that less acetate was being produced, or that it was partly consumed. Model predictions showed that, in this scenario, more acetyl-CoA is converted to pyruvate through PFOR. However, despite higher rates, final biomass concentrations in batch cultivations were slightly lower in the presence of acetate (10 or 25 mM). Our model showed an increase of flux through acetate:CoA ligase (*acs*; EC 6.2.1.1) in the presence of acetate (10 mM), which catalyses acetate assimilation in *Escherichia coli* [84]. This reaction consumes ATP, which would explain the lower biomass concentrations observed. We also observed a higher flux of acetate and acetyl-CoA towards butyrate via butanoyl-CoA:acetate CoA-transferase activity (catalysed by Pct). Batch cultivation experiments did not show a noticeable increase in butyrate concentration when acetate was present, rather lower. Therefore, we hypothesise that, *in vivo*, most acetate consumed is assimilated via acetate:CoA ligase instead of Pct. This deviation to the model seems reasonable since dFBA tends to maximize for biomass and this could be achieved by sending a higher flux of acetate to butyrate instead of assimilating it, saving ATP.

Overall, this work shows the advantages of using a model-driven approach to gain insights into the metabolism of microorganisms. The new findings fill in knowledge gaps and unravel key metabolic features of *A. neopropionicum*. As a result, this study means a step forward to further exploit this species as a cell factory for propionate production both, in mono-culture or in co-cultivation from sustainable feedstocks, e.g., syngas, as recently stated by Moreira et al. [15]. Additionally, *A. neopropionicum* can act as an intermediate species to extend the range of products from propionate to longer odd-chain carboxylic acids.

## Conclusions

In this study, we have constructed iANEO_SB607, the first GEM of *A. neopropionicum*. Combining experimental data with a manual curation of the annotated genome and a comprehensive network reconstruction, we have gained insight into the central carbon and energetic metabolism of this microorganism. The model predicted the metabolic capabilities of *A. neopropionicum* with high accuracy, which allowed us to investigate with detail the enzymatic routes and cofactors involved in the fermentation of ethanol to propionate. Our analysis showed that *A. neopropionicum* produces propionate via propionate-CoA:lactoyl-CoA transferase, the characteristic enzyme of the acrylate pathway. Our *in silico* analysis revealed, for the first time in this microorganism, the presence of the electron-bifurcating Nfn complex. This model provides the basis to explore the capabilities of *A. neopropionicum* as microbial platform for the production of propionate from dilute ethanol as substrate. While beyond the scope of this study, the construction of this model signifies a step closer towards the development of multi-species models that describe syngas-fermenting co-cultures comprised of acetogens with ethanol-consuming propionigenic bacteria. Follow-up studies that integrate, e.g., omics analyses with data from steady-state fermentations should help improve this GEM.

## Availability of data and materials

The supplementary information and the data generated during the current study are available in the following https://gitlab.com/wurssb/Modelling/Anaerotignum_neopropionicum

## Author’s contributions

SBV and IPO conceived and designed the study, and drafted the manuscript. SBV constructed the model and performed model predictions and data analysis. IPO performed the experiments and data analysis. TV contributed to model construction. MSD and PS conceived, designed and supervised the research. DS and VMdS acquired project funding, conceived and supervised the research. All authors reviewed and edited the study. All authors read and approved the final manuscript.

## Acknowledgements

We thank Bart Nijsse for his valuable contribution on the re-annotation of *A*.*neopropionicum*’s genome. We also thank Martijn Diender for the discussions and useful comments on the article.

## Additional Files

Additional File 1 — Genome-scale metabolic modelling to decipher ethanol metabolism via the acrylate pathway in the propionate-producer *Anaerotignum neopropionicum*.

Additional file 1 contains additional figures and tables.

