## Supplementary File for "Genome-scale metabolic modelling enables deciphering ethanol metabolism via the acrylate pathway in the propionate-producer Anaerotignum neopropionicum"

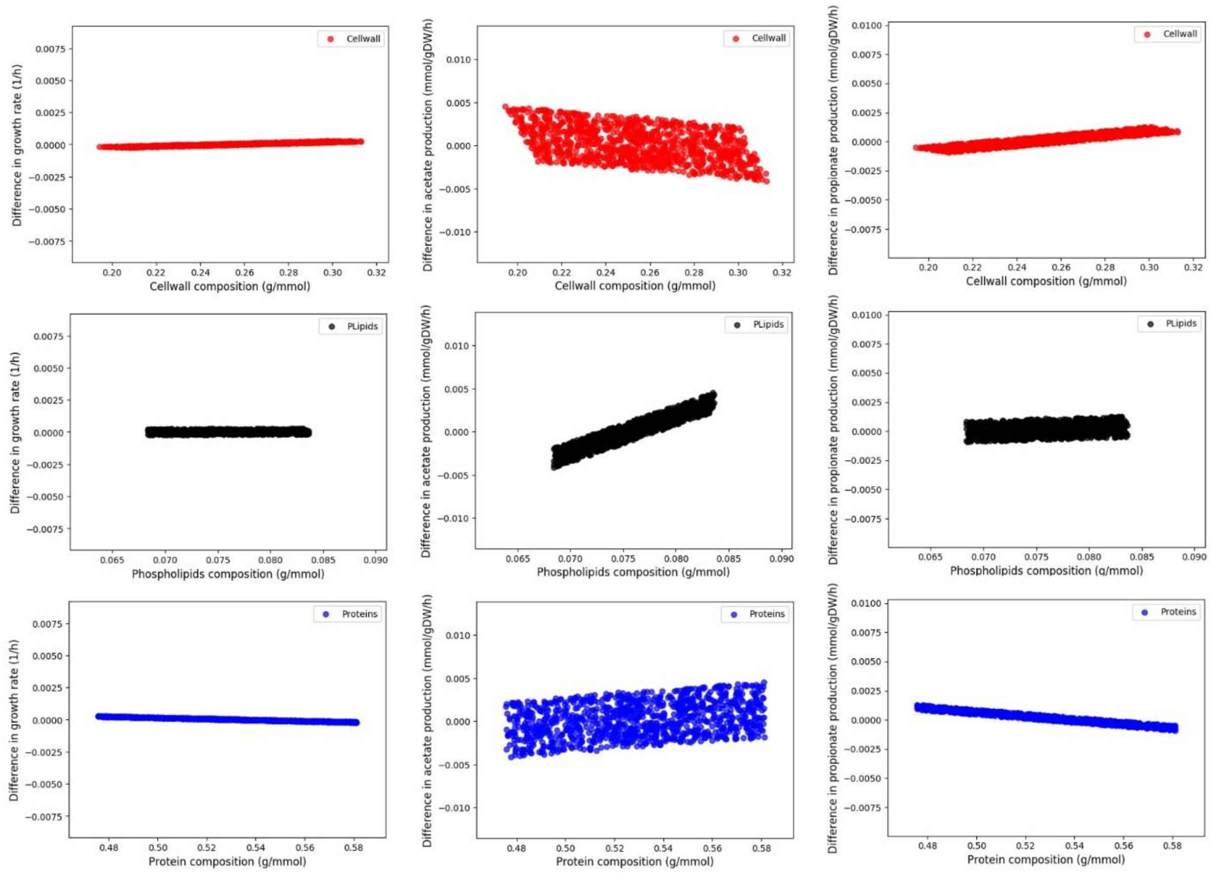


**Figure S1.** Sensitivity analysis of the biomass reaction. Effect of varying the composition of main biomass building blocks on the growth rate and product formation. Growth rate and product formation are represented as the difference between the values obtained when the new biomass reaction is defined as objective function and the values obtained when the original biomass synthesis reaction is defined as objective function.

**Table S1.** Parameters used to simulate batch fermentations through dFBA. Column 'Source' indicates whether the parameter was constrained based on the experimental value (considering standard deviation error) or by model fitting.

| Parameter | Symbol | Value | Units | Source |
|---|---|---|---|---|
| Ethanol fermentation (Fig. 3) | | | | |
| Initial biomass concentration | $X_0$ | 0.0055 | g $L^{-1}$ | Exp. Value |
| Initial Ethanol concentration | $S_{eth,0}$ | 23 | mM | Exp. Value |
| Initial Acetate concentration | $S_{ac,0}$ | 0.15 | mM | Exp. Value |
| Initial propionate concentration | $S_{prop,0}$ | 0.09 | mM | Exp. Value |
| Initial butyrate concentration | $S_{but,0}$ | 0 | mM | Exp. Value |
| Initial propanol concentration | $S_{ppoh,0}$ | 0 | mM | Exp. Value |
| Maximum growth rate | $\mu_{max}$ | 0.082 | $h^{-1}$ | Exp. Value |
| Michaelis-Menten constant for ethanol | $Km_{,etoh}$ | 11 | mM | Fitting (~ exp. value) |
| Maximum ethanol uptake | $q_{etoh,max}$ | 36.5 | mM | Exp. Value |
| Ethanol + acetate fermentation (Fig. 4) | | | | |
| Initial biomass concentration | $X_0$ | 0.008 | g $L^{-1}$ | Fitting (~ exp. value) |
| Initial Ethanol concentration | $S_{eth,0}$ | 23.3 | mM | Exp. Value |
| Initial Acetate concentration | $S_{ac,0}$ | 9 | mM | Exp. Value |
| Initial propionate concentration | $S_{prop,0}$ | 0.1 | mM | Exp. Value |
| Initial butyrate concentration | $S_{but,0}$ | 0 | mM | Exp. Value |
| Initial propanol concentration | $S_{ppoh,0}$ | 0 | mM | Exp. Value |
| Maximum growth rate | $\mu_{max}$ | 0.098 | $h^{-1}$ | Exp. Value |
| Michaelis-Menten constant for ethanol | $Km_{,etoh}$ | 14 | mM | Fitting (~ exp. value) |
| Maximum ethanol uptake | $q_{etoh,max}$ | 43.6 | mM | Exp. Value |

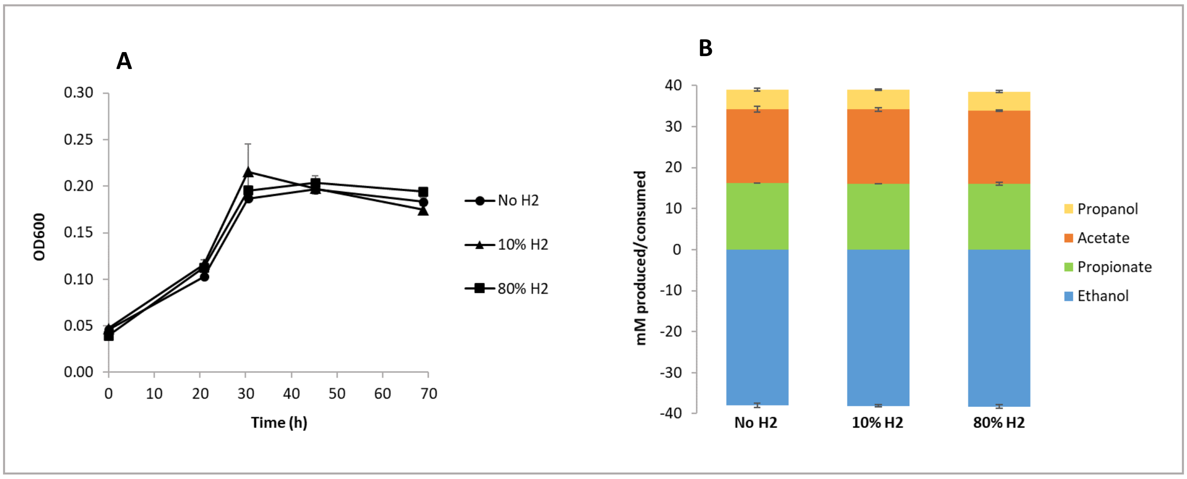


**Figure S2.** Effect of $H_2$ on ethanol-growing cultures of *Anaerotignum neopropionicum*. **A)** Cell growth profiles, determined by optical density at 600 nm (OD600). **B)** End products and ethanol consumed at the end of batch fermentations. Error bars indicate the standard deviation of biological triplicates.

**Table S2.** Fermentation balance of batch cultures of *A. neopropionicum* cultivated on different substrates. Note that $CO_2$, present in the headspace of bottles, is consumed but not included in this table. iBut: isobutyrate; iVal: isovalerate. Traces are concentrations < 0.2 mM. The hyphen symbol indicates undetected products. ND indicates not determined.

| Substrate[a] | Substrate consumed (mM) | Products (mM) | | | | | | |
|---|---|---|---|---|---|---|---|---|
| | | Propionate | Acetate | Propanol | Butyrate | Lactate | iBut | iVal |
| Ethanol | 22.5 ± 0.6 | 9.5 ± 0.1 | 8.6 ± 0.0 | 1.3 ± 0.0 | 0.9 ± 0.0 | - | traces | traces |
| Ethanol + Acetate[b] | 23.0 ± 0.5 | 11.3 ± 0.2 | 16.7 ± 0.3 | 1.2 ± 0.1 | 0.5 ± 0.0 | - | traces | traces |
| Ethanol + Acetate[c] | 23.0 ± 0.3 | 12.5 ± 0.2 | 28.7 ± 0.4 | 0.8 ± 0.1 | 0.3 ± 0.0 | - | traces | traces |
| DL-Lactate | 13.5 ± 1.2 | 8.7 ± 0.5 | 6.2 ± 0.4 | - | - | ND | - | - |
| Glucose | 13.1 ± 1.4 | 11.3 ± 0.2 | 8.7 ± 0.1 | traces | 1.4 ± 0.0 | 0.9 ± 0.0 | traces | traces |
| Xylose | 18.5 ± 0.9 | 15.0 ± 0.7 | 11.0 ± 0.3 | traces | 1.1 ± 0.1 | 2.9 ± 0.2 | traces | traces |

[a] Except for acetate, all substrates were added at a concentration of 25 mM.
[b] Concentration of acetate was 10 mM. The concentration of substrate consumed corresponds to ethanol.
[c] Concentration of acetate was 25 mM. The concentration of substrate consumed corresponds to ethanol.
